# Irradiated Mammary Spheroids Elucidate Mechanisms of Macrophage-Mediated Breast Cancer Recurrence

**DOI:** 10.1101/2022.07.24.501296

**Authors:** Benjamin C. Hacker, Erica J. Lin, Dana C. Herman, Alyssa M. Questell, Shannon E. Martello, Rebecca J. Hedges, Anesha J. Walker, Marjan Rafat

## Abstract

While most patients with triple negative breast cancer receive radiotherapy to improve outcomes, a significant subset of patients continue to experience recurrence. Macrophage infiltration into radiation-damaged sites has been shown to promote breast cancer recurrence in pre-clinical models. However, the mechanisms that drive recurrence are unknown. Here, we developed a novel spheroid model to evaluate macrophage-mediated tumor cell recruitment. We first characterized infiltrating macrophage phenotypes into irradiated mammary tissue to inform our model. We then established spheroids consisting of fibroblasts isolated from mouse mammary glands. We observed that tumor cell motility toward irradiated spheroids was enhanced in the presence of a 2:1 ratio of pro-healing:pro-inflammatory macrophages. We also measured a significant increase in interleukin 6 (IL-6) secretion after irradiation both *in vivo* and in our model. This secretion increased tumor cell invasiveness, and invasion was mitigated by neutralizing IL-6. Taken together, our work suggests that interactions between infiltrating macrophages and damaged stromal cells facilitates breast cancer recurrence through IL-6 signaling.

## Introduction

Almost 300,000 American women will be diagnosed with breast cancer in 2022^1^, and approximately 15% of those patients will have triple negative breast cancer (TNBC), a particularly aggressive subtype. To treat this disease, patients elect to undergo chemotherapy, surgery, and radiation therapy (RT)^2^. RT typically produces positive outcomes for a majority of patients^3,4^. However, an emerging body of literature implicates RT in contributing to cancer recurrence. Up to 20% of patients will experience locoregional recurrence after RT, and this may be correlated to patient immune status^2,5–7^. Lymphopenia, or low lymphocyte count, has been identified as a risk factor for worse clinical outcomes in breast cancer patients^8^. Chemotherapy and RT can cause lymphopenia and immune dysfunction^9^, and lymphopenia is correlated with lower overall survival after therapy^10^. This suggests that a significant number of TNBC patients may benefit from additions to their therapeutic regime.

The impact of normal tissue radiation damage on cancer recurrence is currently understudied. Previous pre-clinical studies have linked radiation to tumor spread. For example, irradiation of mammary stroma induced breast cancer in injected epithelial cells^11^. However, radiation damage alone does not dictate recurrence. Radiation-induced metastasis of TNBC cells to the lung was shown to be facilitated by macrophages^12^. Pre-irradiation of tumor beds led to myeloid cell infiltration and then tumor cell growth due to matrix degradation^13^. More recently, macrophages were shown to promote tumor cell recruitment following normal tissue radiation damage under lymphopenic conditions^5^. However, the contribution of stromal-macrophage interactions to breast cancer recurrence following normal tissue radiation remains unknown.

Spheroid models provide an avenue to determine mechanisms of recurrence,^14^ and their increased biological relevance compared to monolayer studies allows for insights into cell movement, direct cell-cell interactions, and cell-extracellular matrix (ECM) interactions^15–17^. Spheroid models complement *in vivo* models, and they serve as a crucial tool to design well-controlled and robust studies. Here, we developed a novel primary fibroblast spheroid model to evaluate the effect of radiation damage on stromal-macrophage interactions that drive recurrence. We first characterized *in vivo* macrophage infiltration to inform our model parameters. We then used our spheroid model to determine the contributions of irradiated normal tissue and macrophage infiltration to TNBC recurrence.

## Results and Discussion

### In vivo characterization of infiltrating macrophages and cytokine secretion resulting from radiation damage

We characterized the phenotypes of macrophages that infiltrated irradiated mouse mammary fat pads (MFPs) in a model of radiation-induced recurrence (**Figure 1A**)^5^. We defined macrophage (CD45^+^CD11b^+^F4/80^+^) phenotypes as follows: as cells expressing high expression of iNOS, CD86, CD64, or MHCII for pro-inflammatory M1 macrophages and IL4Rα, Arg-1, CD206, or IL-10 for pro-healing M2 macrophages^18–20^. We observed no increase in M1 macrophage infiltration 10 days post-RT. However, a significant increase in M2 macrophage infiltration was observed with tumor cell recruitment (**Figure 1B, Figures S1C, D**). In TNBC patients, higher numbers of tumor associated macrophages typically with an M2 phenotype are correlated with poorer prognosis^21^. Expression of CD163, an M2 marker, in breast tumors was linked to increased likelihood of early distant recurrence and decreased overall survival^22^. Additionally, it has been shown that M2 macrophages play a role in radiation-induced recurrence in other cancers. For example, expression of CD163 in primary head and neck cancer tumors is associated with a higher likelihood of recurrence after RT^23^.

**Figure 1.**
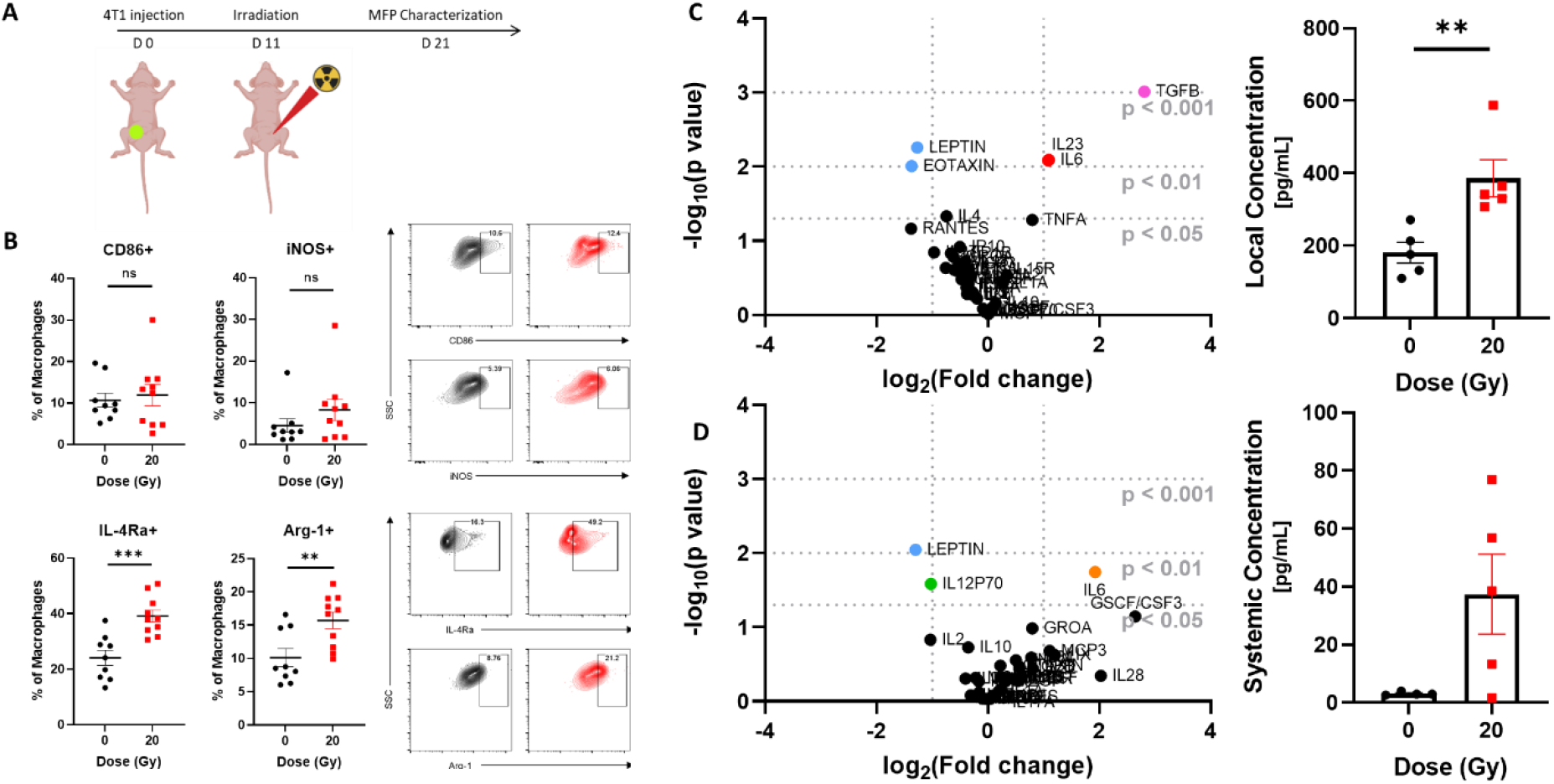
M2 macrophage infiltration and secreted inflammatory cytokines are increased following *in vivo* RT of mammary tissue. **A**. Experimental schematic of *in vivo* normal tissue irradiation studies. **B**. Flow cytometry characterization of M1 and M2 macrophage infiltration into MFPs 10 days post-RT. **C**. Volcano plot of relative change in cytokines in irradiated MFPs. Differences were calculated by normalizing cytokine median fluorescence intensity values 10 days post-RT to values from non-irradiated CD8+ T cell-reduced mice with no tumor. **D**. Volcano plot of relative change in cytokines in serum samples. Error bars show standard error of the mean with **p<0.01 and ***p<0.001 as determined by a two-tailed unpaired t-test.

Because communication between damaged tissue and immune cells is essential to the wound healing response, we evaluated how local and systemic cytokine expression influenced tumor cell recruitment in Balb/C mice with antibody-reduced CD8+ T cells to model lymphopenic patients. Cytokine secretion 10 days post-RT from MFP homogenates and mouse serum was evaluated using a Luminex multiplex immunoassay. Median fluorescence intensity (MFI) values were normalized to those obtained from mice without tumor cell infiltration or RT. MFPs with tumor cell infiltration post-RT exhibited greater than a 2-fold increase in interleukin 6 (IL-6) and IL-23, and more than a 3-fold increase in transforming growth factor beta (TGF-β) relative to a mouse without tumor cell infiltration (**Figure 1C**). Additionally, increases in IL-6 expression were observed systemically (**Figure 1D**). Increased levels of IL-6 in serum are associated with higher risk of early recurrence and bone metastasis^24^. Although IL-6 is typically known as a pro-inflammatory cytokine^25^, it has also been shown to promote TNBC epithelial-mesenchymal transition, progression, cancer stemness, and M2 macrophage polarization^26^. The role of M2 secreted IL-6 as a response to neuroinflammation may be similar to the response of M2 macrophages of a radiation-damaged normal tissue microenvironment^27^. Together, these data indicate that IL-6 influences the recruitment of circulating tumor cells *in vivo*.

### Primary fibroblast spheroid model

The stroma in the mammary gland is made up of a heterogeneous collection of fibroblasts. We evaluated the contributions of cells derived from the mammary stromal vascular fraction (SVF) to tumor cell recruitment. Within 16 hours of plating in U-bottom low-adhesion plates, cells formed spheroids of approximately 300 μm in diameter (**Figure 2A**), a size that is below the threshold where hypoxia and necrosis may occur^14,28^. Surface markers of primary SVF cells were evaluated. The isolated cells expressed high and consistent levels of platelet-derived growth factor receptor beta (PDGFRβ), Podoplanin, and CD90.2/Thy1.2 (**Figure 2B, Figure S2A**). Expression of these markers have been shown in both healthy mammary glands and in mammary glands containing cancer associated fibroblasts^29,30^. There was negligible expression of immune, epithelial, and endothelial markers (**Figure 2B, Figure S2B**), implying that this model is composed primarily of fibroblasts. Fibroblasts produce ECM components, and radiation induces fibrosis, which results in excess ECM production^31,32^. Previous studies have incorporated primary fibroblasts into spheroids^33^; however, we are the first to study the effects of irradiation on spheroid behavior.

**Figure 2.**
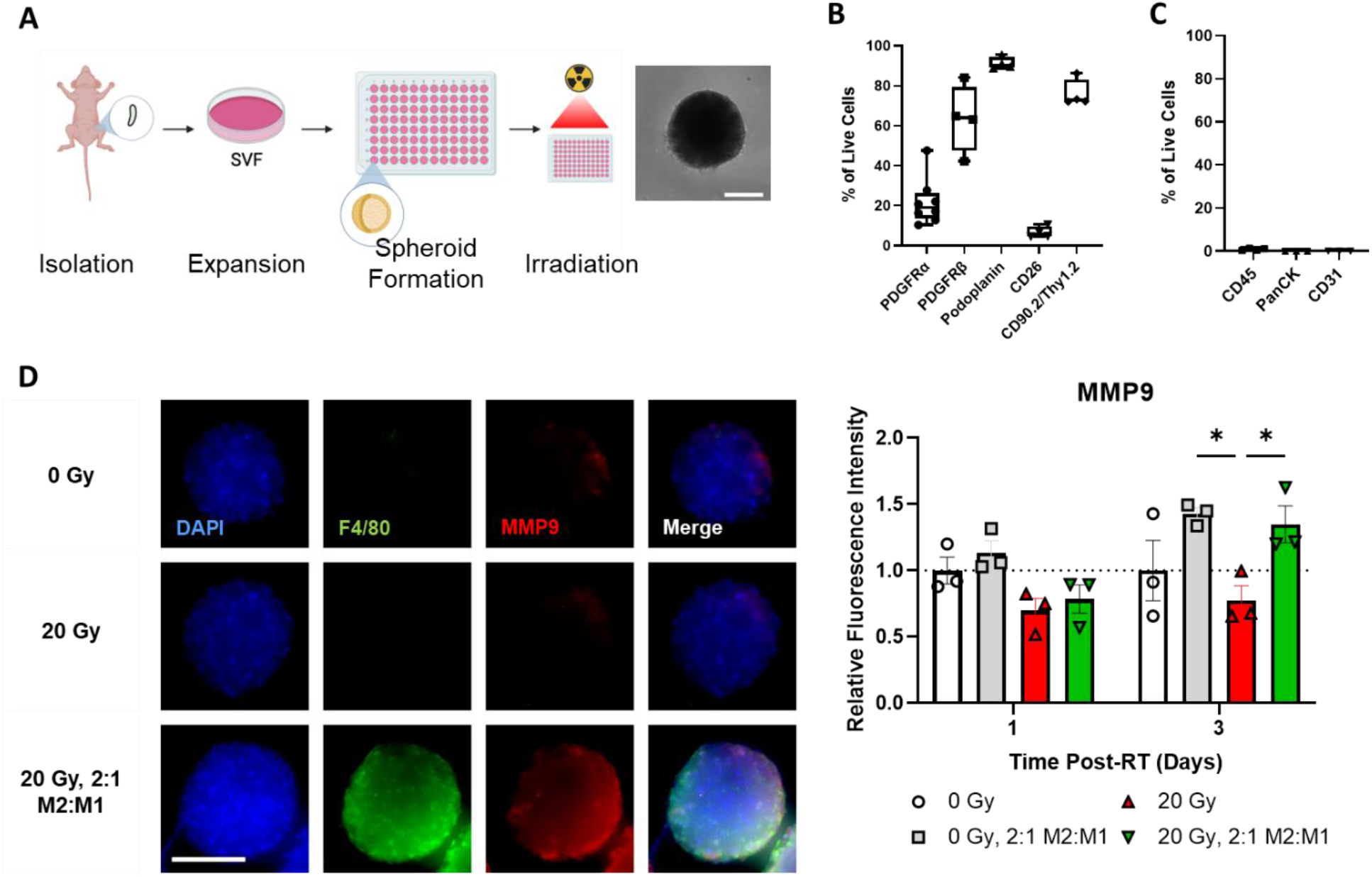
Design of primary mammary spheroids. **A**. Experimental schematic of stromal vascular fraction (SVF) isolation from mouse MFPs, expansion, and irradiation. Brightfield image of representative spheroid. Scale bar is 100 μm. **B**. SVF cells are composed of cells of fibroblast lineage based on flow cytometry characterization. Data is presented as a box and whiskers plot, with the minimum, maximum, first quartile, median, and third quartile shown. **C**. SVF cells show negligible expression of immune, epithelial, and endothelial cells. **D**. Spheroid macrophage (macs) co-culture facilitates enhanced secretion of MMP9. Representative images of immunofluorescence staining of nuclei (blue), F4/80+ macrophages (green), and MMP9 (red) in spheroids 3 days post-RT. MMP9 fluorescence intensity was quantified using Fiji and normalized to 0 Gy for each time point (n = 3). Scale bars are 200 μm. Error bars show standard error of the mean with *p<0.05 as determined by ANOVA.

### The irradiated microenvironment facilitates TNBC cell recruitment through direct spheroid-macrophage interactions

Radiation induces significant infiltration of macrophages in normal mammary tissue^5^, and excess macrophages can cause tumor growth, metastasis, and recurrence^5,34^. We added primary mouse bone marrow derived macrophages (BMDMs) to our spheroid model to study how macrophage infiltration influences tumor cell behavior. Other studies have modeled macrophage infiltration using multiple tissue types^35^. Based on the results of our *in vivo* studies, we added macrophages at a proportion of 15% of the overall cells. M0, M1, and M2 macrophages made up 85%, 5%, and 10% of the overall macrophage population, respectively. To replicate *in vivo* observations of macrophage infiltration, macrophages were added immediately after irradiation (**Figure S2C**,**D**). Using scanning electron microscopy, we observed adhesion and infiltration of macrophages and 4T1s into spheroids after irradiation (**Figure S2E**). Macrophage co-culture and irradiation did not cause any significant differences in spheroid size or aspect ratio (**Figure S2F**). We confirmed infiltration of M1 and M2 macrophages into spheroids after 24 hours (**Figure S2G**).

Matrix metalloproteinase 9 (MMP9) stimulates primary TNBC tumor growth, angiogenesis, and metastasis^36^, but the contribution of local secretion of MMP9 in the irradiated microenvironment to recurrence is relatively unexplored. We observed enhanced MMP9 expression in spheroids co-cultured with macrophages 3 days following RT (**Figure 2D**). This suggests that upon infiltration, macrophages facilitate ECM degradation that may further prime the microenvironment for 4T1 invasion.

We also confirmed that 4T1 murine TNBC cells infiltrated into spheroids (**Figure S2H**). We tracked individual cell infiltration via live cell imaging (**Figure 3A**). When 4T1 cells were co-cultured with macrophages in irradiated spheroids, they showed significantly higher motility toward the center of the spheroid. GFP intensity in the inner core of the spheroid was normalized to the overall GFP intensity of the spheroid after 24 hours of infiltration (**Figure 3B**). In the case of 2:1 M2:M1 macrophages, interior GFP intensity significantly increased, indicating that 4T1 cells were more invasive when cultured in spheroids that recapitulate a microenvironment that has been shown to promote tumor cell recruitment *in vivo*^5^. Additionally, GFP intensity was measured as a function of radial distance from the center (**Figure 3C**). After 72 hours of co-culture, GFP intensity closer to the interior of the spheroid was significantly increased in irradiated spheroids with macrophages relative to both the stromal only irradiated and unirradiated controls. These results show that direct interactions between macrophages and the irradiated stroma facilitate 4T1 recruitment. Macrophages maintained relevant levels of expression of M1 (CD86) markers and M2 (IL4Rα) markers for up to 4 days within the co-culture (**Figure S2I**).

**Figure 3.**
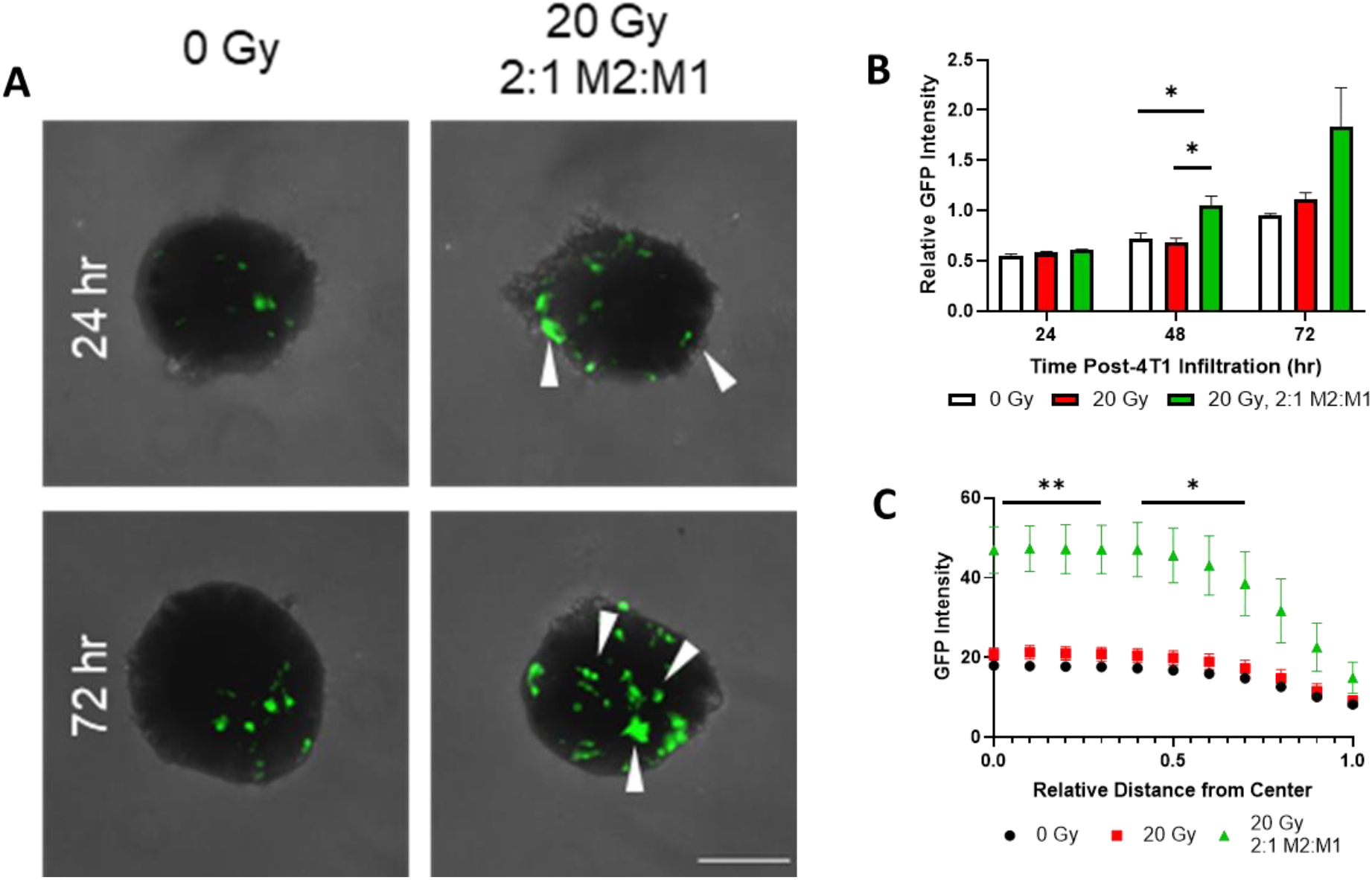
Cell infiltration into spheroids is enhanced by co-culture of irradiated stroma and infiltrating macrophages. **A**. GFP-labeled 4T1 infiltration was tracked at 24- and 72-hours following co-culture with spheroids. Infiltration was enhanced in the presence of 2:1 M2:M1 macrophages co-cultured with irradiated spheroids. **B**. GFP signal from 4T1s is enhanced in the interior of the spheroid over time (n=3-4 biological replicates). Interior spheroid GFP intensity was normalized to the total GFP intensity within the entire spheroid at 24 hours of co-culture. **C**. GFP intensity increases with macrophage co-culture as a function of radial distance at 72 hours. Error bars show standard error of the mean with *p <0.05 and **p <0.01 as determined by a two-tailed unpaired t-test. White arrows indicate cell infiltration. Scale bars are 200 μm.

### IL-6 secretion drives 4T1 cell invasiveness

We isolated conditioned media (CM) from co-cultures of macrophages and irradiated spheroids. We characterized IL-6 secretion from spheroids as a function of irradiation and macrophage infiltration to determine if *in vivo* cytokine secretion is replicated in our model. We observed that infiltration of macrophages (2:1 M2:M1) resulted in a significantly higher secretion of IL-6 relative to spheroids with no macrophage infiltration (**Figure 4A**). Additionally, when macrophages were co-cultured with unirradiated spheroids, we observed no significant changes in IL-6 secretion. Interestingly, the co-culture of M0 macrophages with irradiated spheroids did not change IL-6 secretion (**Figure S3E**). These observations establish the importance of maximizing biological relevance by incorporating multiple macrophage phenotypes into spheroid models. The impact of IL-6 on 4T1 invasiveness was then investigated using a transwell assay. 4T1 invasiveness was significantly higher when incubated with CM from irradiated spheroids with macrophages relative to CM collected from spheroids with no macrophages (**Figure 4B**), and neutralization of IL-6 in CM resulted in nearly fourfold reduction in invasiveness of 4T1 cells. Invasion was also significantly lower in CM from unirradiated spheroids co-cultured with macrophages. Previous studies have shown that neutralization of IL-6 reduces invasiveness of 4T1 cells and that macrophage-derived IL-6 contributes to tumor cell migratory ability^37,38^. However, we are the first to show the dependence of 4T1 invasiveness on IL-6 secreted from stromal-macrophage interactions.

**Figure 4.**
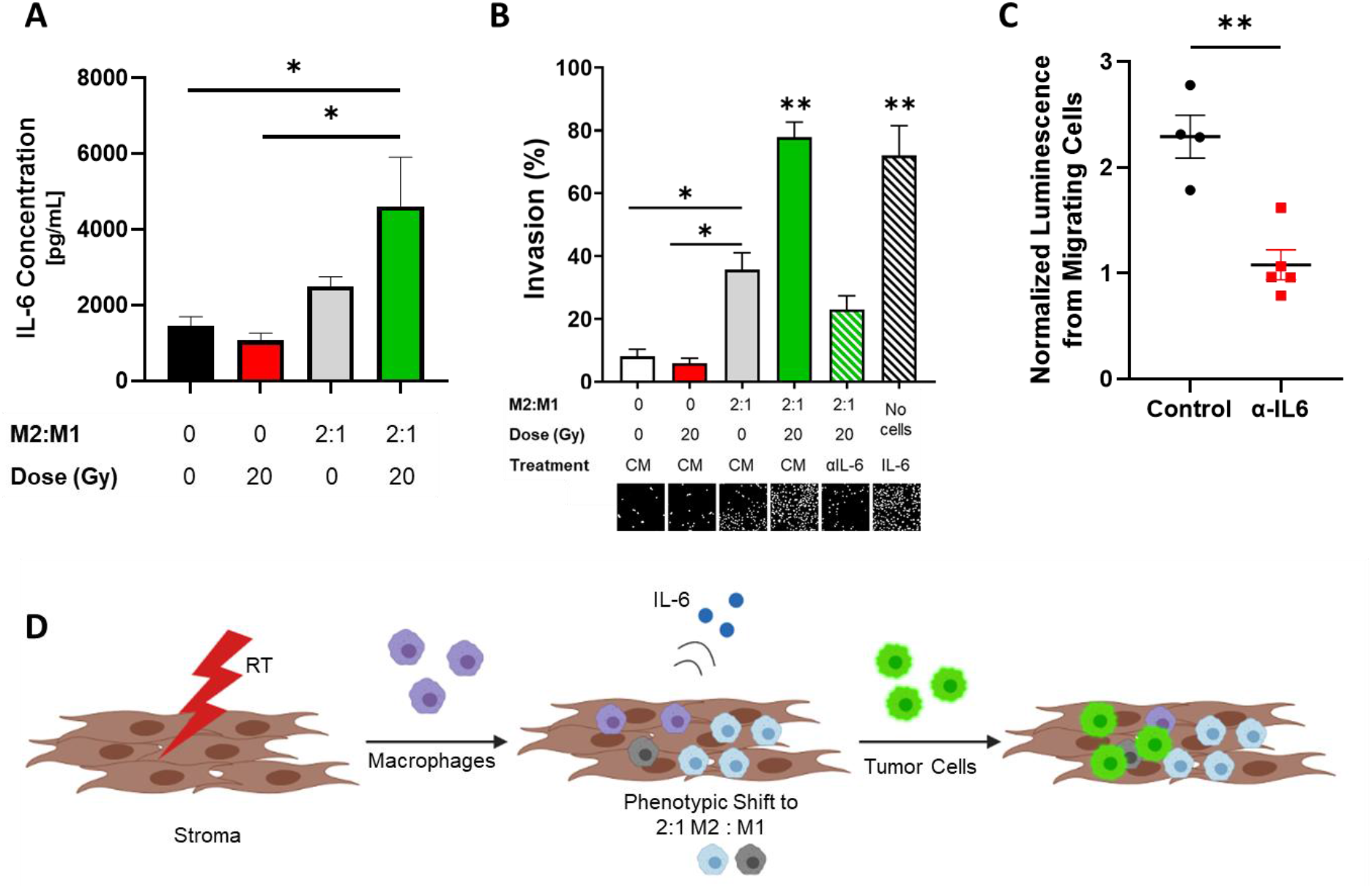
IL-6 secretion drives tumor cell recruitment and invasion post-RT. **A**. Spheroid conditioned media (CM) incubated for 24 hours after RT promotes IL-6 secretion when cultured with a 2:1 ratio of M2:M1 macrophages (green). IL-6 secretion is lower in the absence of spheroid RT (black), with spheroid RT and no macrophage infiltration (red), or with macrophage co-culture without RT (gray). **B**. IL-6 secreted from RT spheroids co-cultured with macrophages promotes 4T1 cell invasion. IL-6 neutralization (αIL-6, striped green) in CM of irradiated spheroids with macrophage co-culture diminished invasion. Error bars show standard error of the mean with *p<0.05 and **p<0.01 as determined by ANOVA analysis. 20 Gy (solid green) and IL-6 spiked media (striped black) are significant to all other conditions. **C**. IL-6 neutralization *in vivo* downregulates 4T1 cell invasion after irradiation. Luminescence from migrating cells was normalized to the unirradiated control for each treatment. Error bars show standard error of the mean with **p<0.01 as determined by unpaired, 2-tailed t-test. **D**. Proposed mechanism of 4T1 tumor cell infiltration following normal tissue RT.

We then investigated the impact of IL-6 on tumor cell recruitment *in vivo*. IL-6 was systemically neutralized 6 hours before irradiation of contralateral MFPs in mice with orthotopic luciferase-labeled 4T1 tumors, and bioluminescence imaging (BLI) was used to quantify the luminescent signal from recruited tumor cells in MFPs. For each treatment, BLI was normalized to the non-irradiated control. Systemic depletion of IL-6 significantly reduced BLI levels after RT (p<0.01; **Figure 4C**), indicating inhibition of tumor cell recruitment. Taken together, these data suggest that the interaction between tissue damaged from RT and M2 and M1 macrophages causes secretion of IL-6 that in turn recruits 4T1 cells (**Figure 4D**). IL-6 depletion was not observed to have significant off-target effects, including primary tumor growth and tumor metastasis, suggesting that this mechanism may be more specific to local recurrence (**Figures S3F-K**). Myeloid-derived IL-6 has been reported in 4T1 progression and metastasis^38,39^. In the clinic, IL-6 inhibitors have been largely unsuccessful at preventing progression of primary disease, showing little efficacy in multiple myeloma, renal cell carcinoma, and prostate cancer^40^. Inhibition of IL-6R has been shown to hinder TNBC metastasis in a pre-clinical model^41^. However, there is limited knowledge of the clinical efficacy of IL-6 and IL-6R inhibitors in primary breast cancer progression and recurrence^42^. This is the first report of IL-6 driving 4T1 invasiveness post-RT in the context of local recurrence. This work highlights the importance of using 3D spheroid models to elucidate mechanisms of cancer recurrence. Our results indicate that monitoring serum IL-6 levels may provide prognostic and therapeutic value in improving outcomes for TNBC patients vulnerable to recurrence.

## Methods

### Cell Lines

Luciferase-labeled and GFP expressing 4T1 mouse mammary carcinoma cells were obtained from Dr. Laura L. Bronsart (Stanford University). All cells were cultured at 37°C and 5% CO_2_. 4T1 cells were cultured in RPMI-1640 (Gibco) supplemented with 10% heat-inactivated fetal bovine serum and antibiotics (100 U/mL penicillin and 100 mg/mL streptomycin). Cells were used within three passages before injection into mice.

### Orthotopic Tumor Studies

Animal studies were performed in accordance with institutional guidelines and animal protocols approved by the Vanderbilt University Institutional Animal Care and Use Committee. Tumor inoculation was performed by injecting 5 × 10^4^ 4T1 cells in a volume of 50 μL of sterile PBS into the number four inguinal right MFPs of 8-to 10-week old female Nu/Nu or Balb/C mice (Charles River Laboratories). In CD8+ T cell reduction experiments, 0.5 mg anti-CD8a (2.43; BioXCell) was injected intraperitoneally every 5 days starting from the day of inoculation^5^. Control mice were injected with 0.5 mg rat IgG2b isotype control (LTF-2; BioXCell) using the same dosing schedule. In IL-6 depletion experiments, 0.5 mg anti-IL6 (BioXCell) was injected intraperitoneally starting 6 hours before irradiation, then every 3 days afterward. Control mice were injected with 0.5 mg IgG1 isotype control antibody (BioXCell). Tumor length and width were measured using digital calipers (Fisher Scientific) beginning one week after tumor inoculation. Tumor volume was calculated as follows: Volume = (L_1_^2^ x L_2_)/2, where L_1_ is the smaller diameter of the tumor, and L_2_ is the larger diameter^5^.

### Luminex Multiplex Cytokine Assay

To assess cytokine profiles at the local site of infiltration, MFPs were harvested and homogenized in 20 mM Tris HCl (pH 7.5) buffer with 0.5% Tween 20, 150 mM NaCl, and protease inhibitor, centrifuged for 10 min at 4°C, and supernatant was stored at -80°C^43^. Protein content was measured using bicinchoninic acid protein assay (ThermoFisher). For evaluation of systemic cytokine signaling, blood samples were collected via retro-orbital bleed, allowed to clot at room temperature for 30 minutes, then centrifuged for 10 minutes. Serum was then recovered and stored at -80°C. Samples were processed at the Stanford Human Immune Monitoring Center using a mouse 39-plex Affymetrix kit.

### Primary Spheroid Generation

Spheroids were generated from primary cells obtained from the SVF in mouse MFPs using previously published techniques^33^. Briefly, MFPs were harvested and minced with a blade and digested for 40 minutes in a solution of PBS with 20 ug/mL liberase, and antibiotics (100 U/mL penicillin and 100 mg/mL streptomycin). Tissue was flushed through a 100 μm filter and plated in 10 cm dishes in DMEM/20% BCS. Upon confluence, cells were plated in low adhesion U-bottom plates at a density of 10,000 cells/spheroid. Spheroids were formed within 24 hours of passaging.

### Radiation

Radiation was delivered by two methods. For *in vivo* RT, mice were anesthetized by administering isoflurane and irradiated to 20 Gy using a 300 kVp cabinet x-ray system filtered with 0.5 mm Cu. The mice were shielded using a Cerrobend jig with apertures 1 cm wide and 1.5 cm long to expose normal MFPs. Transmission through the 2 cm thick shield was less than 1%. For *in vitro* experiments, spheroids were irradiated in U-bottom low adhesion plates to 20 Gy using a Cesium source.

### Flow Cytometry

MFPs were harvested and minced in a solution of 1% FBS in PBS. They were then placed in a 2 mg/mL solution of Collagenase II (Sigma) and incubated for 30 minutes in a water bath at 37°C. The digestion was inactivated by adding 1% FBS in PBS. The digested tissue was then passed through a 100 μm filter to generate a single cell suspension. Cells were stained with the Aqua fixable viability stain (ThermoFisher) and FC receptors were blocked with CD16/32 (Biolegend) simultaneously with other cell surface markers for 20 minutes at 4°C. After staining, cells were rinsed with PBS and fixed with 1% neutral buffered formalin in saline for at least 20 minutes at 4°C. Intracellular stains were performed using an intracellular permeabilization buffer (ThermoFisher). Fixed cells were rinsed with PBS for 5 mins, rinsed with permeabilization buffer for five minutes, then incubated with antibodies diluted in permeabilization buffer for thirty minutes at room temperature in the dark. Cells were then rinsed with permeabilization buffer and resuspended in PBS. Flow cytometry was performed on a four-laser Amnis CellStream machine (Luminex), and FlowJo software was used for analysis. Compensations were obtained by using compensation beads (ThermoFisher). The following antibody clones were used for analysis: CD45 (30-F11), CD11b (M1/70), F4/80 (BM8), MHCII (M5/114.15.2), iNOS (CXNFT), CD64 (X54-5/7.1), CD86 (GL1), CD206 (C068C2), IL-4Rα (I015F8), IL-10 (JES5-16E3), and Arg-1 (A1exF5). For staining of primary SVF cells, cells were passaged and stained using the same staining protocol. The following antibody clones were used for analysis: PDGFRα (APA5), PDGFRβ (APB5), Podoplanin (8.1.1), CD26 (H194-112), and CD90.2/Thy1.2 (53-2.1).

### Spheroid Embedding, Sectioning, and Immunofluorescence

Spheroids were stained using previously published methods^44,45^. Tissues were carefully transferred to microcentrifuge tubes using wide bore pipet tips. Spheroids were rinsed in PBS and fixed in 10% Neutral Buffered Formalin overnight at 4°C. They were then rinsed in PBS and incubated in 30% sucrose overnight at 4°C. They were subsequently incubated 2-4 hours in 1:1 mixture of 30% sucrose-PBS and OCT, transferred to a mold with OCT, and stored at -80°C before cryosectioning. For immunofluorescence, slides were incubated in PBS to remove OCT and permeabilized with 0.1% Triton-X 100 in PBS for 10 minutes. Sections (15 μm) were then rinsed with PBS and blocked with 1% bovine serum albumin (BSA) in PBS with 10% normal goat serum (NGS) for 1 hour at room temperature in humid chambers. Sections were incubated with primary antibody diluted in 1% BSA in PBS overnight in humid chambers at 4°C. After rinsing with PBS, sections were incubated with secondary antibody diluted in 1% BSA in PBS for 2 hours in humid chambers. Then, after additional PBS rinses, sections were incubated with phalloidin (ThermoFisher) for 1 hour. Coverslips were mounted onto slides using NucBlue mounting media (ThermoFisher) and allowed to cure overnight before imaging on a fluorescence microscope (Leica DMi8). Antibodies and stains used for immunofluorescence studies include iNOS (Abcam), CD206 (Abcam), collagen IV (Abcam), MMP 9 (Abcam), F4/80 (Invitrogen) and phalloidin (ThermoFisher).

For whole mount staining, spheroids were transferred to PBS after fixation. They were blocked and permeabilized in buffer containing 1% BSA, 1% DMSO, 1% Triton-X 100, and 1% NGS for 1 hour at room temperature on an orbital shaker. After blocking, they were incubated with primary antibodies for 72 hours at 4°C. After primary antibody incubation, spheroids were rinsed 5 times, then incubated with secondary antibodies (1:200), Hoechst (2 μM), and phalloidin (Thermofisher) for 24 hours at 4°C. After secondary incubation, spheroids were rinsed 5 times and imaged. Fiji was used to for image analysis.

### Scanning Electron Microscopy (SEM)

Spheroids were fixed using methods previously published^46^. Briefly, spheroids were carefully transferred from culture plates into a microcentrifuge tube using a wide bore pipet. Spheroids were washed in 1x PBS and then fixed in 2% glutaraldehyde, 4% paraformaldehyde, and 0.1 M sodium cacodylate buffer (pH 7.3) for 1 hour at 4°C followed by incubation in 1% glutaraldehyde in PBS for 15 min at room temperature. After fixation, spheroids were washed 3x in PBS then dehydrated in a series of 10 minute incubations of 10%, 25%, 50%, 75%, 90%, and 2x 100% ethanol at room temperature. The spheroids were then further dehydrated in 50% ethanol:hexamethyldisilazane (HMDS) and 100% HMDS, and air dried overnight. They were mounted on carbon backed aluminum stubs, sputter coated with gold for 45 seconds (Cressington), and imaged using a Quanta 250 Environmental Scanning Electron Microscopy at 5 kV.

### Primary Macrophage Isolation and Culture

Bone marrow derived macrophages (BMDMs) were cultured using previously published methods^47–49^. Briefly, macrophages were isolated from the femurs of Nu/Nu mice. Femurs were crushed using a mortar and pestle, and the cell suspension was passed through a 40 μm filter. Red blood cells were lysed with an ACK lysis buffer, and macrophage precursors were plated at densities of 1×10^6^ cells/plate in IMDM supplemented with 10% FBS, antibiotics (100 U/mL penicillin and 100 mg/mL streptomycin), and 10 ng/mL macrophage colony stimulating factor (MCSF) on 10 cm low adhesion plates for 7 days for maturation into macrophages. For polarization, media was spiked with LPS and IFN-γ for 24 hours (M1 macrophages) or IL-4 for 48 hours (M2 macrophages).

### Invasion Assay

Conditioned media (CM) collected from spheroid and macrophage co-cultures was used as chemoattractants in a transwell migration assay (Corning reduced growth factor Matrigel invasion chamber, 8 μm pore size). Spheroids were irradiated to 20 Gy and then co-cultured with 2:1 M2:M1 macrophages. Supernatant was collected after 24 hours of co-culture. 100,000 4T1 cells were seeded into the upper chambers and incubated with CM for 24 hours. Cells that invaded through Matrigel inserts or migrated through uncoated inserts were stained with NucBlue mounting media and nuclei were counted. Invasion was calculated by dividing invasion counts by migration counts.

### Spheroid 4T1 Migration Assay

24 hours after irradiation of spheroids, 100 GFP-labeled 4T1 cells were plated per spheroid. Infiltration was monitored via live cell fluorescence and phase contrast imaging every 24 hours for a total of 72 hours after plating of 4T1 cells. GFP signal was evaluated as a function of radial position via a custom MATLAB (MathWorks) script. Along the radial axis, 1,000 readings of fluorescent intensity were evaluated over every angle of each spheroid. Readings were normalized to the overall length of each radius to generate relative intensity values. For each biological replicate, more than 10 technical replicates were analyzed.

### IL-6 Quantification

IL-6 secretion in CM was measured using an ELISA assay (R&D systems) per manufacturer instructions.

## Statistical Analysis

To determine statistical significance, differences in radiation dose on flow cytometry data and cytokine concentration were analyzed using two-tailed unpaired t-tests. One-Way ANOVA was evaluated when comparing co-culture conditions for GFP intensity, IL-6 secretion, and 4T1 invasiveness. All statistical analysis was performed in GraphPad Prism 9.

## Acknowledgments

The authors thank Dr. Laura L. Bronsart for providing the GFP- and luciferase-labeled 4T1 cells, Dr. Michael Freeman for irradiator use, Dr. Austin Kirschner for the Cerrobend jig, and Dr. Craig L. Duvall for IVIS use. We also thank the Vanderbilt Cell Imaging Shared Resource (CISR) core for SEM imaging and the Translational Pathology Shared Resource (TPSR) for IF sample preparation. Schematics were prepared by using BioRender.

This research was financially supported by NIH grants #R00CA201304, F31CA254311 (B.C. Hacker), S10RR026373 (CISR), and P30CA068485 (TPSR) as well as the Breast Cancer Alliance Young Investigator Grant.

The authors declare no competing financial interests.

Author contributions: M. Rafat and B.C. Hacker conceived the study, designed experiments, and wrote and revised the manuscript. B.C. Hacker, E.J. Lin, D.C. Herman, A.M. Questell, S.E. Martello, R.J. Hedges, and A.J. Walker conducted experiments and processed and analyzed the data supervised by M. Rafat.

## Notes

### Competing Interest Statement

The authors have declared no competing interest.

### Summary of Updates

Updated results/figures and text.

