## Supplementary Figures for "Irradiated Mammary Spheroids Elucidate Mechanisms of Macrophage-Mediated Breast Cancer Recurrence"

Running Title: Spheroids Reveal Cancer Recurrence Mechanisms

Benjamin C. Hacker[1], Erica J. Lin[2], Dana C. Herman[3], Alyssa M. Questell[4], Shannon E. Martello[1], Rebecca J. Hedges[1], Anesha J. Walker[5], Marjan Rafat[1,4,6,*]

[1]Department of Chemical and Biomolecular Engineering, Vanderbilt University, Nashville, TN, USA

[2]Department of Biological Sciences, Vanderbilt University, Nashville, TN, USA

[3]Department of Biochemistry, Vanderbilt University, Nashville, TN, USA

[4]Department of Biomedical Engineering, Vanderbilt University, Nashville, TN, USA

[5]Department of Biology, Tennessee State University, Nashville, TN, USA

[6]Department of Radiation Oncology, Vanderbilt University Medical Center, Nashville, TN, USA

*Author for correspondence: Marjan Rafat, PhD, Engineering and Science Building, Rm. 426, Vanderbilt University, Nashville, TN 37212. Phone: (615) 343-3389, Fax: (615) 343-7951,

### Supplementary Figures

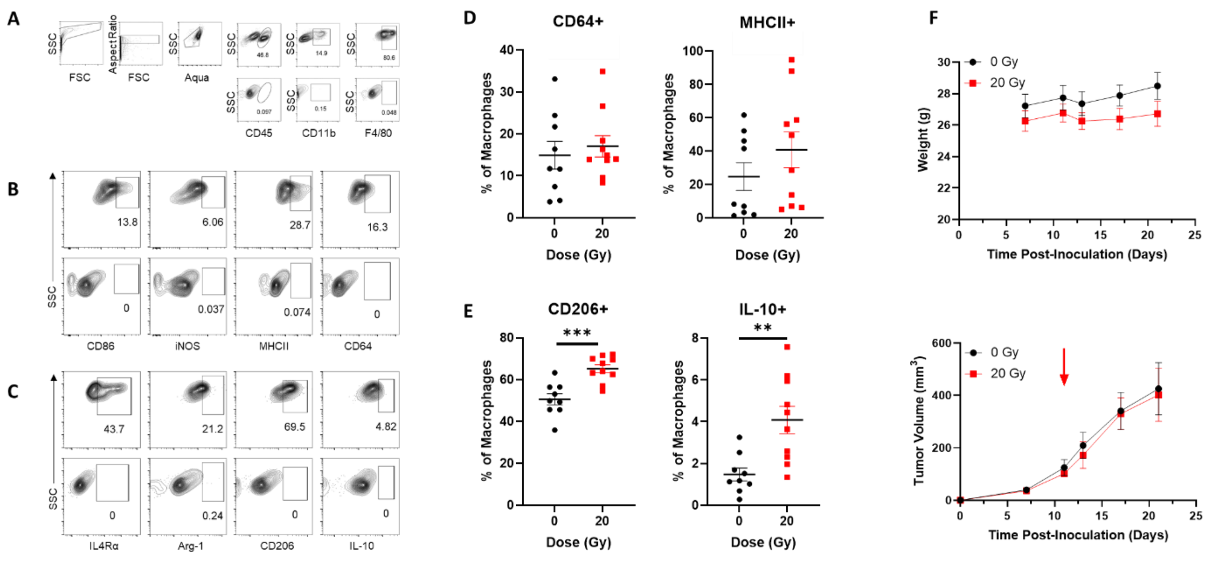


**Supplementary Figure S1. Macrophage infiltration in normal tissues 10 days post-RT A.** Flow cytometry gating strategy for infiltrating macrophages in mammary fat pads (MFPs) with unstained controls. Flow cytometry gating strategy for M1 (**B**) and M2 (**C**) macrophages with unstained controls. Quantification of infiltrating F480+ M1 (**D**) and M2 (**E**) macrophages post-RT. Expression of CD64 and MHCII (M1) and CD206 and IL-10 (M2) were quantified. **F.** Mouse weight and tumor volume curves. Arrow indicates time of irradiation. Data is mean ± standard error of mean. N=9-10 biological replicates per treatment. **p<0.01 and ***p<0.001 as determined by a two-tailed unpaired t-test.

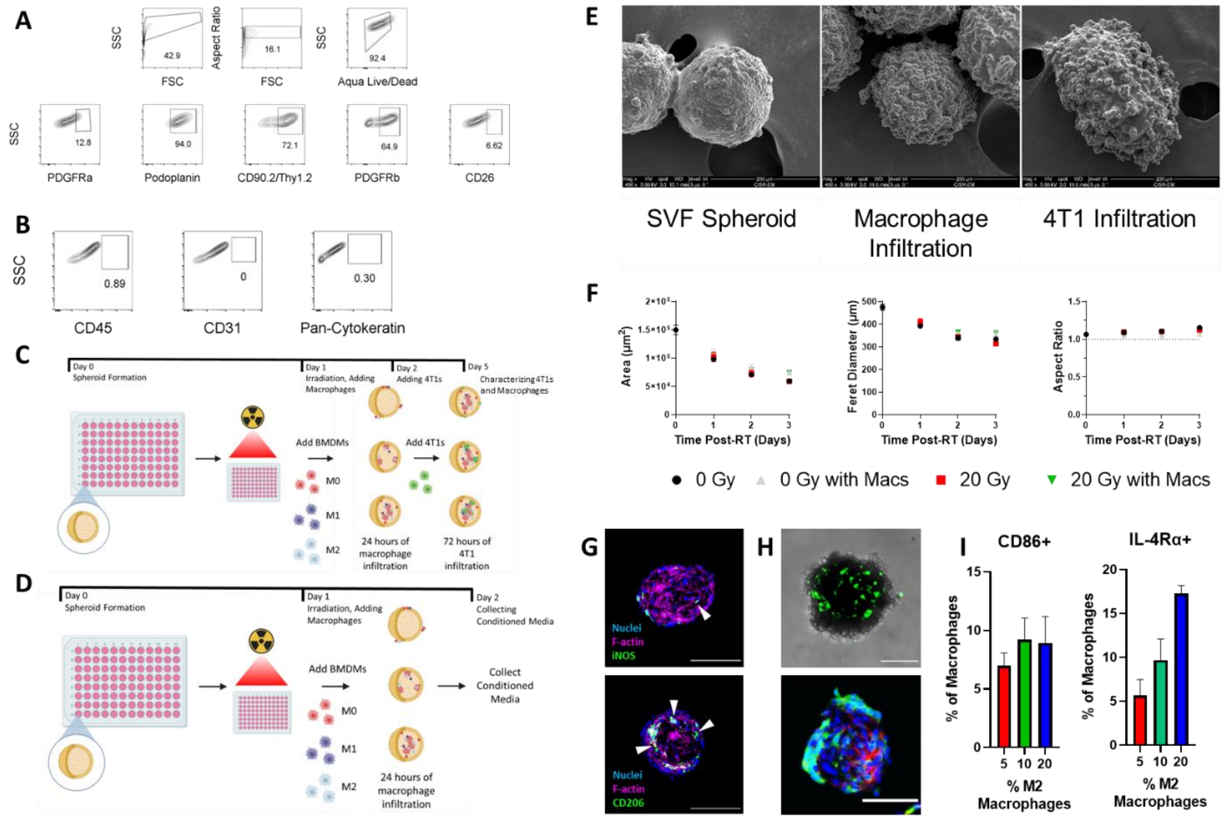


**Supplementary Figure S2. SVF spheroid characterization. A.** Full gating strategy for SVF markers. **B.** Negligible expression of immune cell, endothelial cell, and epithelial cell markers. **C.** Schematic of 4T1 co-culture experimental setup. **D.** Schematic of conditioned media experimental setup. **E.** Scanning electron microscopy images of SVF spheroids alone, with macrophage infiltration, and with 4T1 infiltration. Images were taken at 400x magnification. **F.** Spheroid area (μm$^2$), feret diameter (μm), and aspect ratio were quantified. **G.** M1 (iNOS) and M2 (CD206) macrophages infiltrate into spheroids within 24 hours. F-actin was used to stain the cytoskeleton (magenta). **H.** Confirmation of 4T1 (GFP-labeled) infiltration via live cell imaging. Sections were stained with phalloidin (red) and NucBlue (blue), respectively. Scale bars are 200 μm. **I.** Confirmation of expression of M1 and M2 markers 4 days after co-culture (n=3 biological replicates). For **F** and **I**, data is mean ± standard error of mean, n=3 biological replicates.

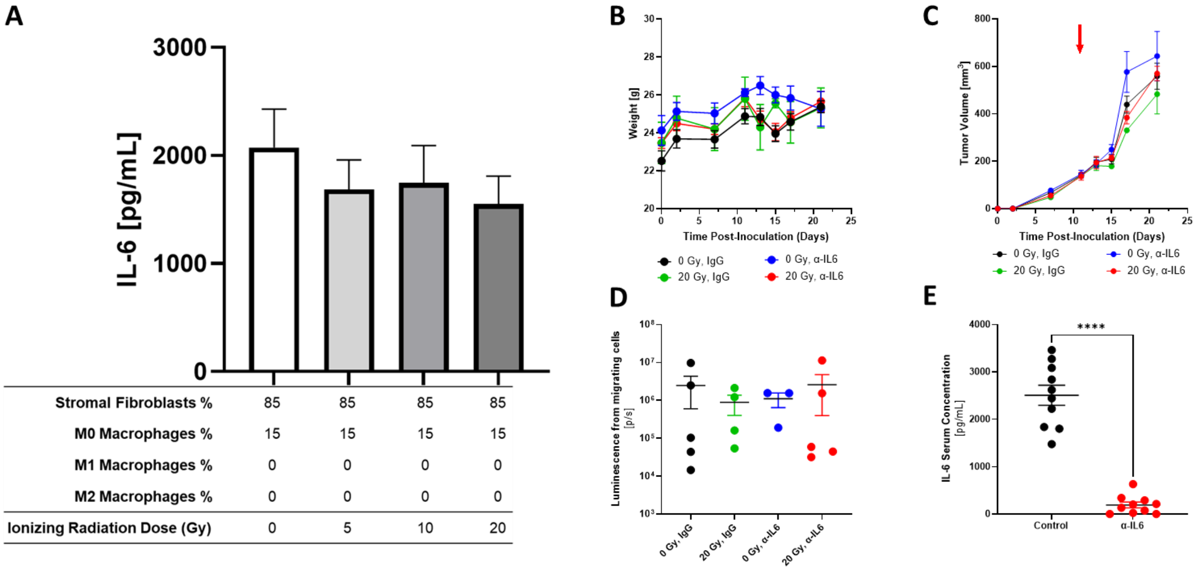


**Supplementary Figure S3. *In vitro* spheroid model co-culture and *in vivo* validation of IL-6 depletion model. A.** IL-6 concentration in co-cultures with M0 macrophages does not change with RT dose. N=3 biological replicates per condition. For *in vivo* experiments, mouse weights (**B**) and tumor curves (**C**) are shown. Arrow indicates time of irradiation. **D.** Metastatic burden in mouse lungs was quantified using BLI. **E.** Systemic IL-6 depletion was confirmed via ELISA. Data is mean ± standard error of mean. N=4-5 biological replicates per treatment. ****p<0.0001, statistical significance determined by a two-tailed unpaired t-test.
